# Horizontal gene transfer drives the emergence of nitrogen fixation in a unicellular *Synechocystis* lineage

**DOI:** 10.64898/2026.04.18.719408

**Authors:** Leonardo Ken Okumura, Mari Banba, Kazuma Uesaka, Shouta Nonoyama, Yuichi Fujita, Shinji Masuda

## Abstract

Nitrogen fixation plays a central role in primary productivity and nitrogen cycling in aquatic ecosystems, yet its distribution among cyanobacterial lineages remains incompletely understood. Biological nitrogen fixation is energetically costly and highly oxygen-sensitive, imposing constraints in oxygenic phototrophs. The unicellular cyanobacterial genus *Synechocystis* has long been regarded as strictly non-diazotrophic. Here, we report that *Synechocystis* sp. LKSZ1 possesses a functional nitrogen fixation system. Comparative genomics revealed that LKSZ1 is distinct from other *Synechocystis* strains and uniquely harbors a complete *nif* gene. Phylogenetic and structural analyses indicate acquisition via horizontal gene transfer from filamentous cyanobacteria. Physiological assays demonstrated photoautotrophic growth under nitrogen-depleted conditions and nitrogenase activity under microoxic to anaerobic conditions. Disruption of *nifK* abolished both growth and activity. These findings show that ecological nitrogen limitation and host compatibility can enable functional integration of horizontally acquired nitrogen fixation.

## Introduction

Biological nitrogen fixation (BNF) serves as a primary biological pathway for converting atmospheric N₂ into bioavailable forms [1–3]. Cyanobacteria are major oxygenic phototrophs capable of BNF [1,4–6], yet diazotrophy is unevenly distributed across cyanobacterial lineages [7,8] and is constrained by the extreme oxygen sensitivity of nitrogenase [1–3]. Consequently, even non-heterocystous cyanobacteria can contribute to nitrogen inputs when physiological and environmental conditions permit nitrogenase activity, with important implications for nutrient dynamics in natural waters [6,9].

The cyanobacterial genus *Synechocystis* has long been regarded as non–nitrogen-fixing and has functioned as a premier model system for photosynthesis, metabolism, and stress responses. Although nitrogen fixation has been reported in a few strains historically assigned to *Synechocystis* [9,10], these early studies relied primarily on morphological and physiological criteria [11], and their phylogenetic status remains unverified by modern phylogenomic standards. Consequently, it remains unclear whether nitrogen-fixing strains attributed to *Synechocystis* truly reside within this genus at the genomic level or instead represent misclassified members of closely related diazotrophic lineages.

Advances in genome-based taxonomy, including species delineation based on average nucleotide identity (ANI) [12] and pangenome analyses that distinguish conserved and flexible gene repertoires [13,14] now provide a framework for revisiting this issue. Phylogenomic studies have further suggested that the common ancestor of several cyanobacterial lineages, including *Synechocystis*, *Microcystis*, *Cyanothece*, and *Crocosphaera*, may have been capable of nitrogen fixation, followed by lineage-specific loss or retention of this trait [8]. Within this evolutionary context, the presence of nitrogen fixation in genomically defined *Synechocystis* necessitates determining its evolutionary origin, namely, whether it represents retention of an ancestral trait or reacquisition via horizontal transfer of a *nif* gene cluster from distantly related cyanobacteria. To date, no study has addressed this question by integrating pangenome structure, genome plasticity, and phylogenetic analyses of *nif* genes in a genome-confirmed *Synechocystis* lineage.

Here, we characterize a unicellular cyanobacterial strain, LKSZ1, previously isolated from a freshwater pond on our university campus (Fig.1 AB) and assigned to the genus *Synechocystis* [15], which harbors a *nif* gene cluster associated with biological nitrogen fixation. We combine genome-based taxonomy, comparative genomics, phylogenetic analyses of the *nif* gene cluster, and physiological and genetic experiments to show that the strain LKSZ1 represents a *Synechocystis* lineage that has functionally acquired *nif*-dependent nitrogen fixation via horizontal gene transfer (HGT), challenging assumptions about metabolic capabilities in this genus and highlighting HGT as a driver of diversification under nitrogen limitation.

**Figure 1.**
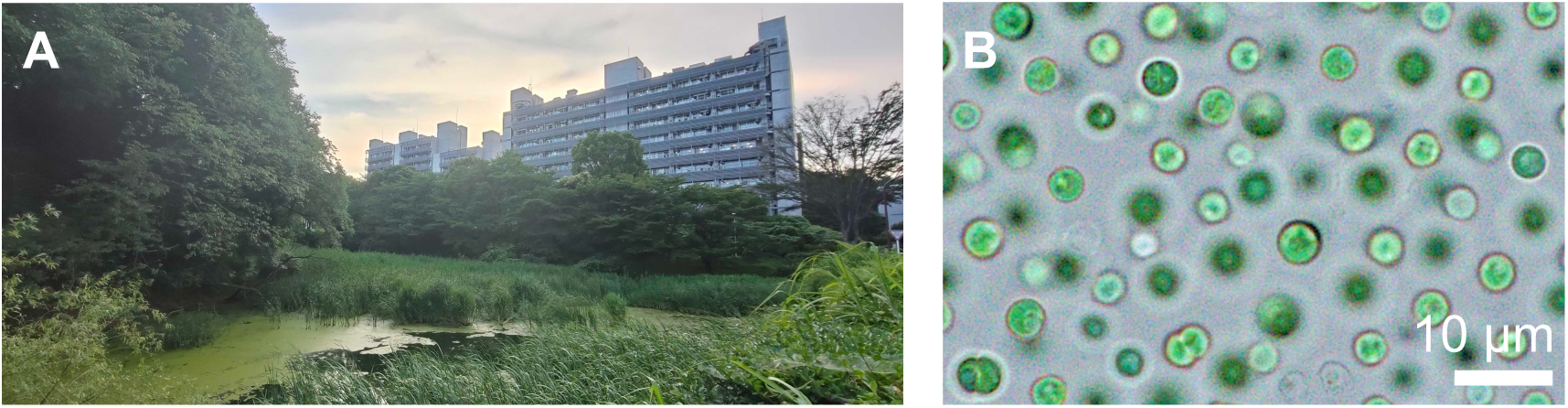
Isolation site and cellular morphology of *Synechocystis* sp. LKSZ1. (A) Freshwater pond (Suzukake Pond, Institute of Science Tokyo, Yokohama, Japan) where *Synechocystis* sp. LKSZ1 was isolated. (B) Light micrograph of strain LKSZ1 grown under standard BG11 conditions. Cells were imaged using a 100× oil-immersion objective lens.

## Results

### Genome-based characterization of *Synechocystis* sp. LKSZ1 and pangenome structure

Figure 2 shows a subtree extracted from the Genome Taxonomy Database (GTDB) representative species phylogeny based on 120 conserved bacterial marker genes. The strain LKSZ1 clusters robustly within the genus *Synechocystis*, distinct from filamentous taxa such as *Thermoleptolyngbya* and forming a broader group with related unicellular diazotrophs like *Crocosphaera*. To establish the genomic position of *Synechocystis* sp. LKSZ1 within the genus *Synechocystis*, comparative analyses were conducted using 32 publicly available *Synechocystis* genomes (Fig. 3). Average nucleotide identity (ANI) analysis revealed that strain LKSZ1 exhibited ∼73% ANI with the model strain *Synechocystis* sp. PCC 6803, significantly below the commonly accepted species boundary of 95% [12]. Similar low ANI values were observed between strain LKSZ1 and other *Synechocystis* strains, indicating that strain LKSZ1 is genomically distinct from previously characterized members of the genus.

**Figure 2.**
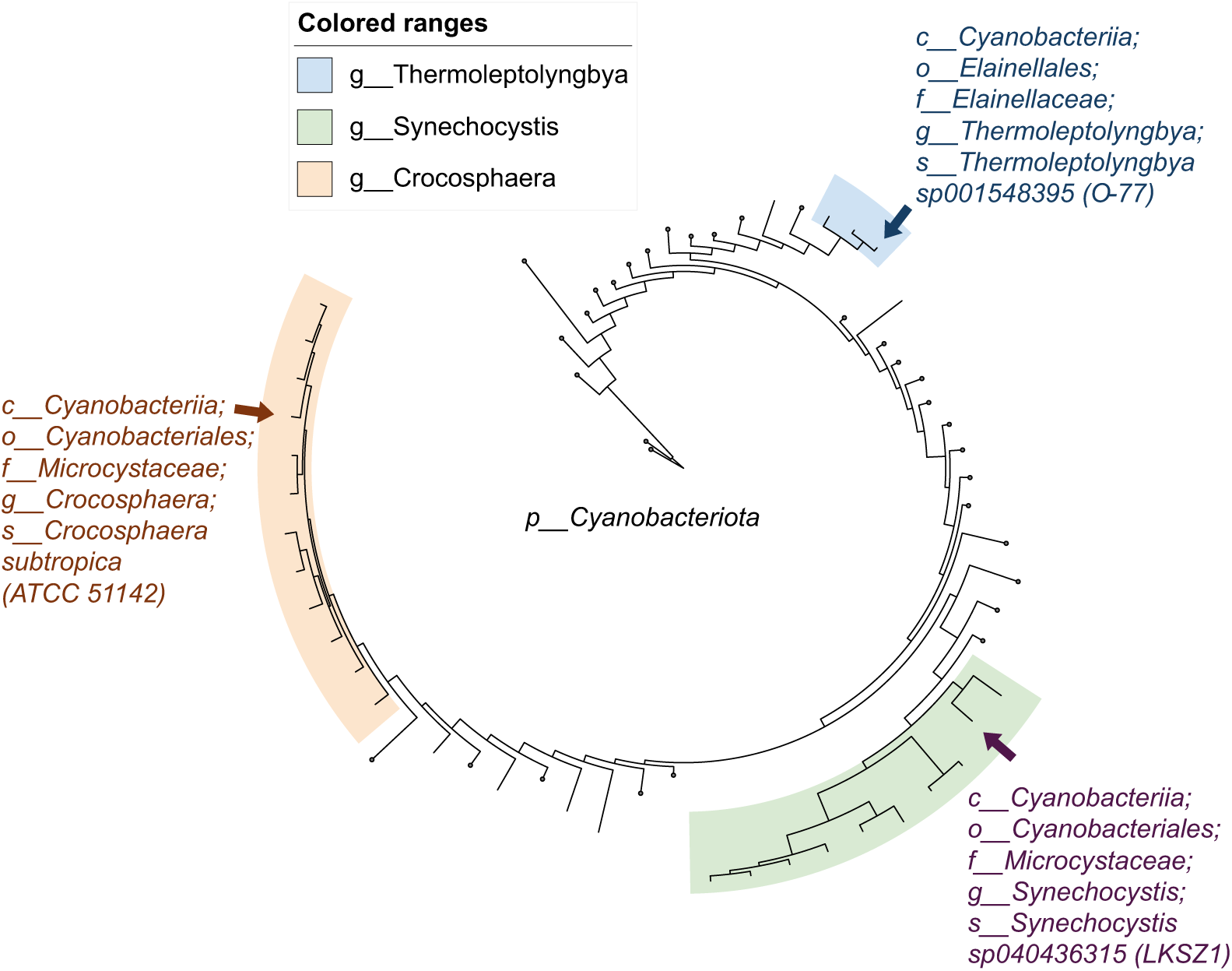
Phylogenomic placement of strain LKSZ1 within the Cyanobacteria. Subtree was extracted from the Genome Taxonomy Database (GTDB) representative species phylogeny based on 120 conserved bacterial marker genes. The tree represents cyanobacterial lineages, with branch lengths proportional to the number of substitutions per site; selected branches are collapsed and shown as circles. *Synechocystis* sp. LKSZ1, *Thermoleptolyngbya* sp. O-77, and *Crocosphaera subtropica* ATCC 51142 are highlighted to illustrate their respective phylogenomic positions. Strain LKSZ1 clusters robustly within the genus *Synechocystis*, distinct from filamentous taxa such as *Thermoleptolyngbya* and forming a broader group with related unicellular diazotrophs like *Crocosphaera*. This genome-scale phylogeny confirms the taxonomic identity of strain LKSZ1 and provides a critical framework for interpreting the phylogenetic incongruence observed in the Nif protein trees (S1 Fig).

**Figure 3.**
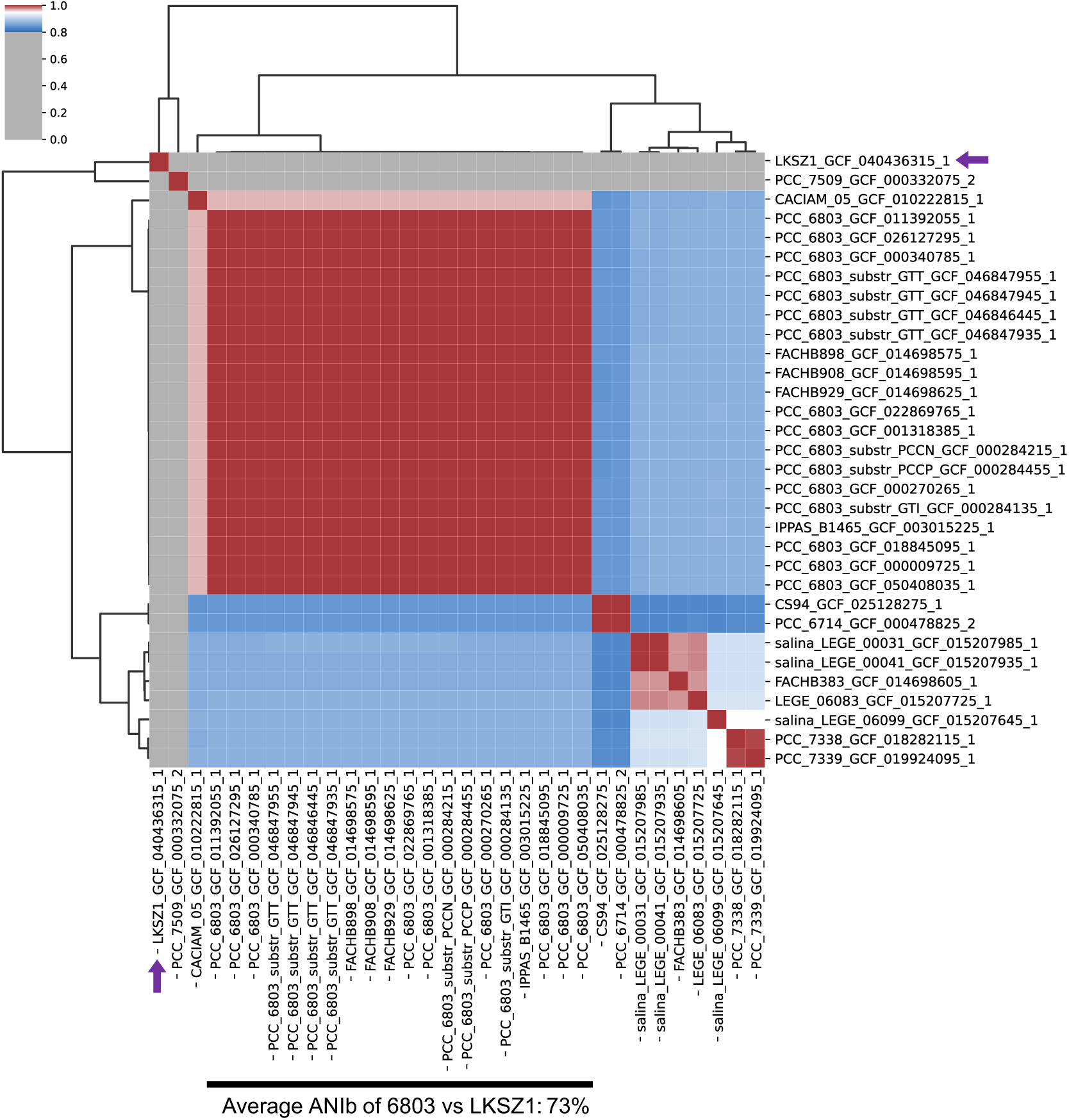
Hierarchical clustered average nucleotide identity (ANI) heatmap among *Synechocystis* genomes. Heatmap showing pairwise ANI values calculated via the ANIb algorithm in pyani-plus (v1.0.0) across 32 *Synechocystis* genomes, including strain LKSZ1. Strain LKSZ1 (purple arrow) forms a distinct cluster separate from *Synechocystis* sp. PCC 6803 and other *Synechocystis* species level classification, with pairwise ANI values below the 95% cutoff (the commonly accepted species boundary (Richter and Rosselló-Móra, 2009)). The dendrogram illustrates hierarchical clustering based on ANI similarity.

Pangenome analysis of the 32 *Synechocystis* genomes identified a total of 3,143 gene families in LKSZ1, which were partitioned into 2,044 persistent (65%), 130 shell (4%), and 969 cloud (31%) gene families (Fig. 4, S1 File). Persistent gene families are conserved across most genomes and represent the core genomic backbone, whereas shell gene families are moderately conserved and variably present among strains, and cloud gene families are rare and often strain-specific. In strain LKSZ1, persistent genes constituted the majority of conserved genomic regions, whereas shell and cloud genes were unevenly distributed along the chromosome.

**Figure 4.**
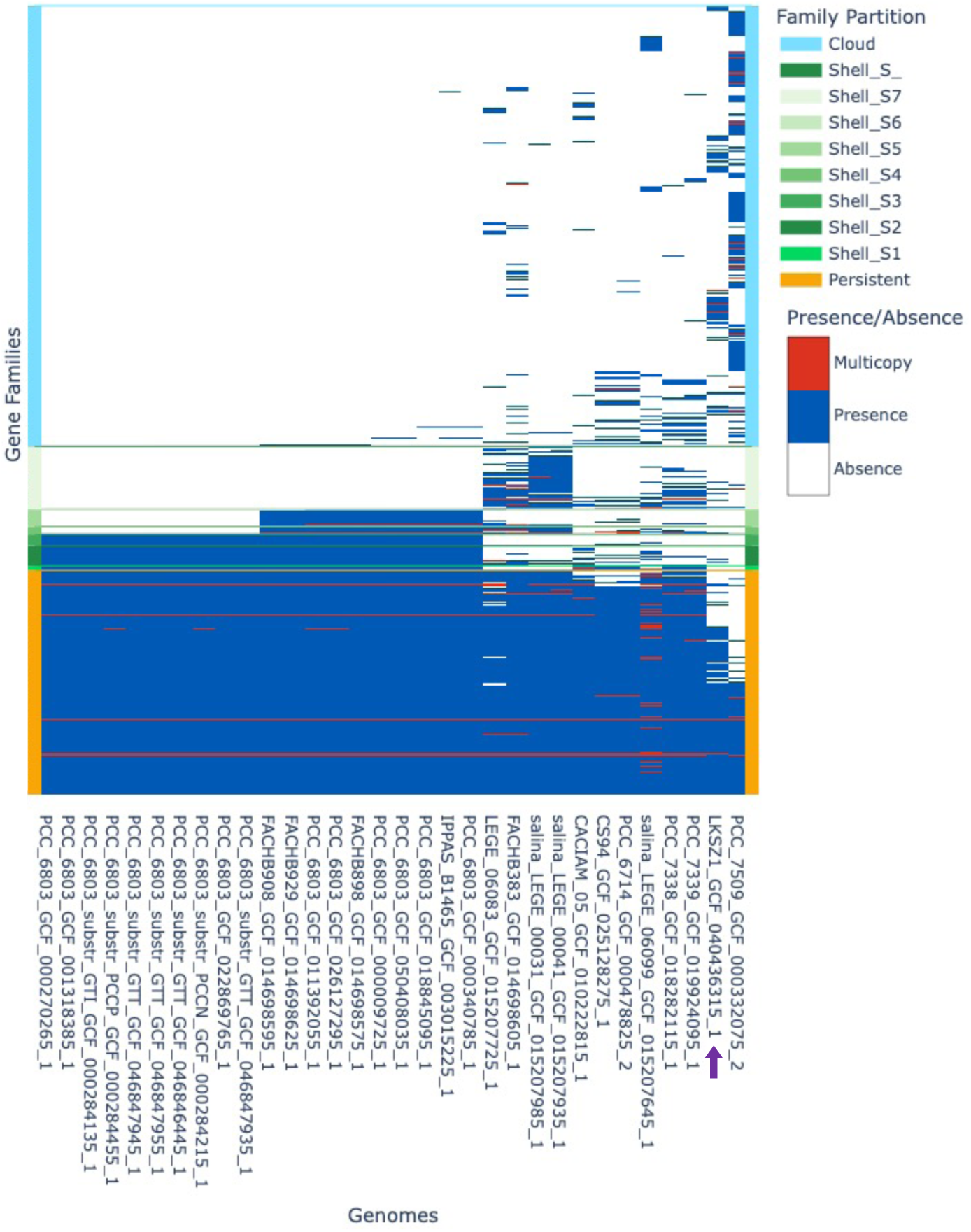
Presence and absence of genes and copy number variation of *Synechocystis* genomes. Tile plot illustrating gene family distribution across 32 *Synechocystis* genomes, including strain LKSZ1 (purple arrow), determined using PPanGGOLiN (v2.2.4). Gene families are partitioned into persistent (orange), shell (green), and cloud (light blue) categories based on the software’s statistical model. Gene family occurrence within each genome is color-coded: red indicates multicopy presence, blue indicates single-copy presence, and white indicates absence. Persistent gene families are highly conserved across genomes (81%), whereas shell and cloud families exhibit variable distribution patterns. The distribution reveals an extensive accessory genome repertoire in strain LKSZ1 relative to other members of the genus.

Statistical genome partitioning further identified 62 regions of genome plasticity (RGPs) in strain LKSZ1, defined as genomic segments enriched in shell and cloud gene families disrupting the persistent genomic backbone (Fig. 5, S1 File). These RGPs were dispersed throughout the chromosome and varied in size and gene content, highlighting a high degree of genomic plasticity within strain LKSZ1 relative to other members of the genus (S1 File).

**Figure 5.**
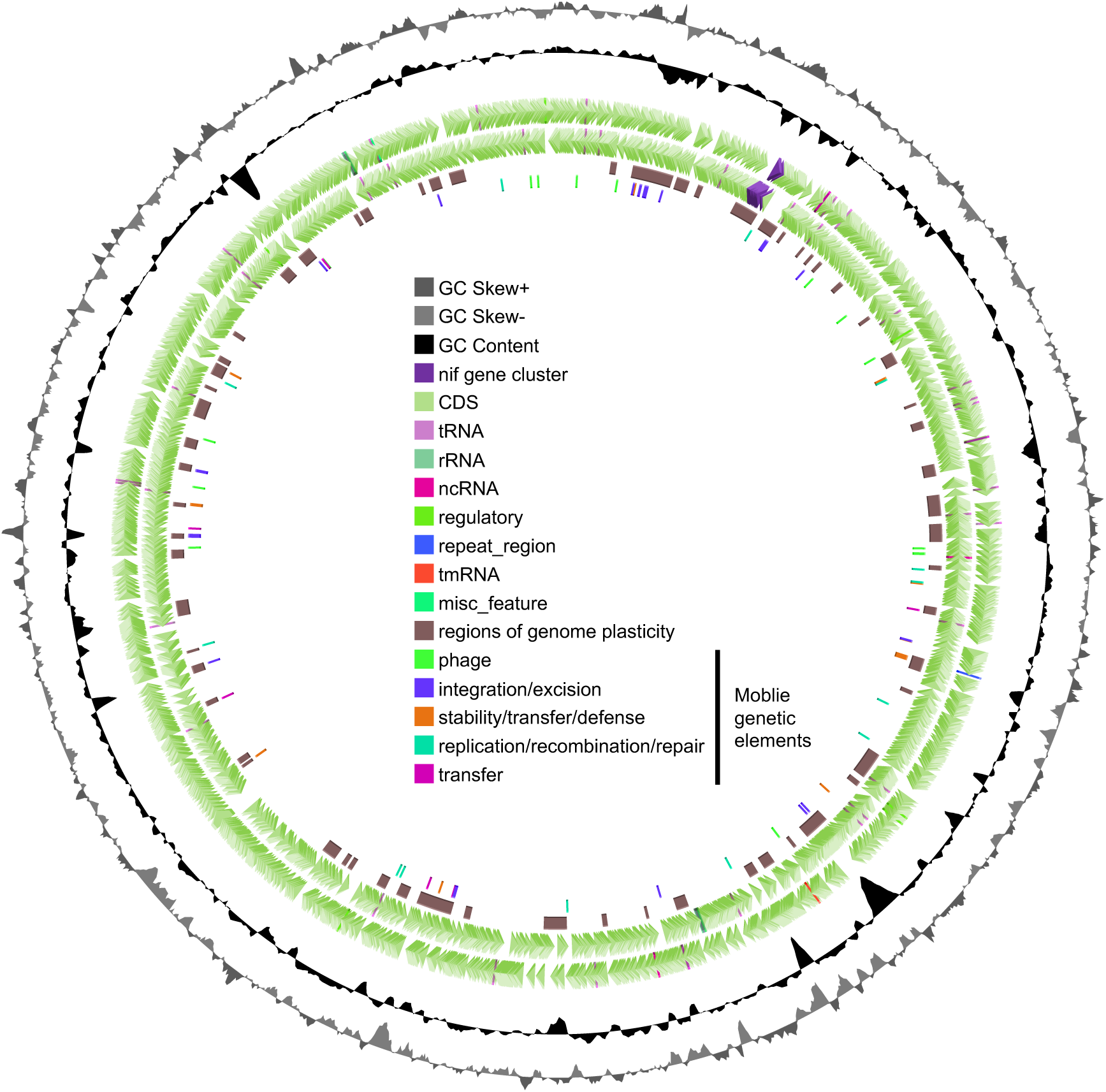
Circular genome map of *Synechocystis* sp. LKSZ1 highlights the *nif* gene cluster and associated genomic features. Circular visualization of the strain LKSZ1 genome generated using Proksee. Tracks are shown from outer to inner: (i) GC skew; (ii) GC content; (iii) coding sequences (CDS) annotated by NCBI RefSeq, with the *nif* gene cluster highlighted in purple; (iv) regions of genome plasticity (RGPs) identified by PPanGGOLiN in brown; and (v) predicted mobile genetic elements identified using mobileOG-db. The *nif* gene cluster is localized within an RGP and coincides with localized deviations in GC content and GC skew relative to the genomic background. These features are consistent with horizontal acquisition and subsequent integration, notwithstanding the absence of identifiable insertion sequences or transposase signatures within or immediately flanking the cluster.

### Identification of a *nif* gene cluster within a genome plasticity region

Genome partitioning identified that one of the RGPs in *Synechocystis* sp. LKSZ1 contained a contiguous ∼29.7 kb cluster of genes associated with biological nitrogen fixation, the *nif* gene cluster (Fig. 6A, B). The *nif* gene cluster in strain LKSZ1 was identified as a continuous genomic region extending from a putative transcriptional regulator gene (*cnfR*) to the molybdate transporter gene *modBC*. The total length of this region was 29,708 bp. Within this interval, genes encoding the core structural components of nitrogenase (*nifH*, *nifD*, and *nifK*), as well as genes required for cofactor biosynthesis (*nifEN* and *nifB*), were tightly clustered. The *nif* gene cluster was organized in the order *nifBSUHDKZEN*, corresponding to the E-type cluster exemplified by *Synechococcus* sp. PCC 7335 [8].

**Figure 6.**
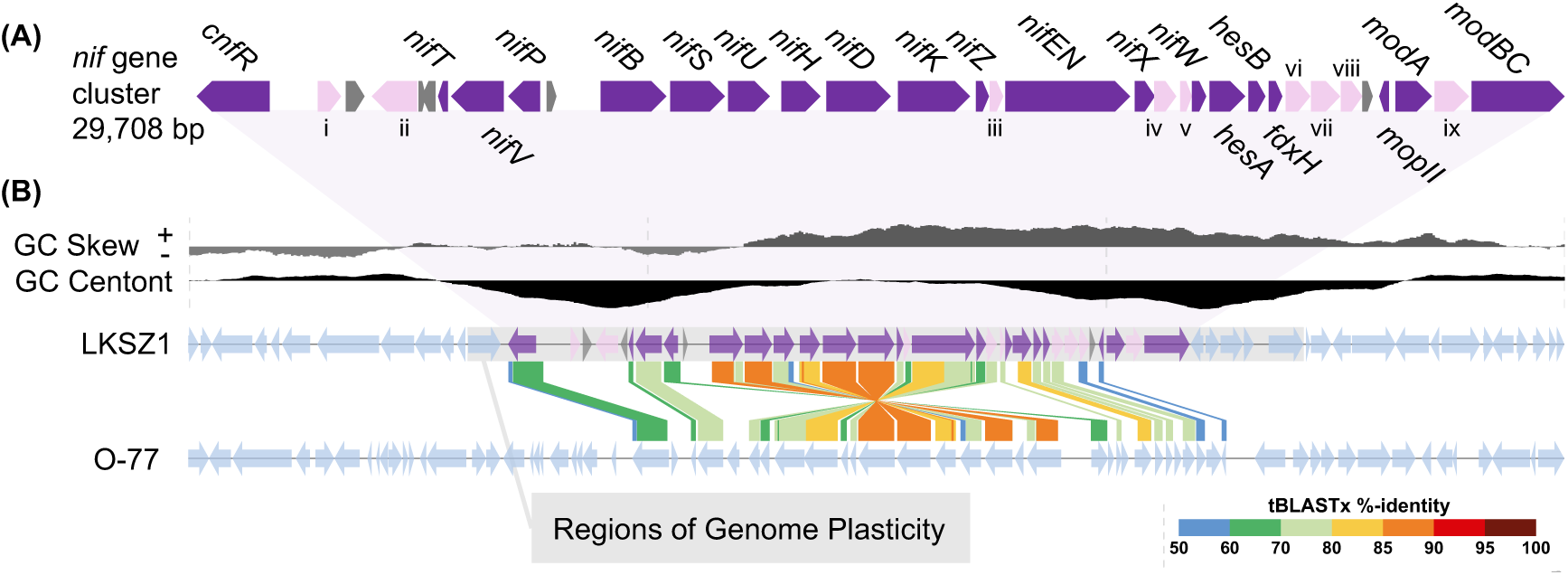
Genomic and structural characterization of the *nif* gene cluster in *Synechocystis* sp. LKSZ1. (A) Organization of the *nif* gene cluster identified in the genome of strain LKSZ1. The cluster spans approximately 29.7 kb and extends from the putative transcriptional regulator gene, *cnfR* to *modBC*. Genes highlighted in purple encode predicted nitrogenase structural components and associated cofactors, including the nitrogenase reductase NifH, nitrogenase α subunit NifD, nitrogenase β subunit NifK, FeMo-cofactor biosynthesis proteins NifEN, NifB, and additional accessory proteins involved in nitrogen fixation. Genes highlighted in pink encode: (i) an aminotransferase class I/II large domain protein, (ii) a ParB/sulfiredoxin domain protein, (iii) a Mo-dependent nitrogenase family protein, (iv) a NifX-associated nitrogen fixation protein, (v) a CCE_0567 family metalloprotein, (vi) flavodoxin, (vii) a cupin-domain protein, (viii) a nitrogenase-associated ArsC/Spx/MgsR family protein, and (ix) a heavy-metal chelation domain protein. Gray arrows represent genes annotated as hypothetical proteins. Arrow direction indicates predicted transcriptional orientation. (B) Genomic context of the *nif* gene cluster in strain LKSZ1. The cluster is located within a region of genome plasticity identified by PPanGGOLiN partitioning. GC content and GC skew (10,000-bp sliding window, 100-bp step) are shown relative to the chromosomal average. Synteny comparison between the 60 kbp region, including the *nif* gene cluster of LKSZ1, and that of *Thermoleptolyngbya* sp. O-77 was performed using tBLASTx-based DiGAlign analysis. Colored links indicate regions of significant sequence similarity at the protein level.

Comparative screening of the 31 other *Synechocystis* genomes revealed that none contained an equivalent *nif* gene cluster or individual nitrogenase structural genes (S2 File). Thus, within the analyzed dataset, strain LKSZ1 was the only member of the genus *Synechocystis* that encoded a complete set of genes required for nitrogen fixation.

### The *nif* gene cluster in *Synechocystis* sp. LKSZ1 was acquired through HGT

To infer the evolutionary affiliation of the *nif* gene cluster in *Synechocystis* sp. LKSZ1, phylogenetic analyses were performed using amino acid sequences of core nitrogenase components and cofactor biosynthesis proteins (NifH, NifD, NifK, and NifEN). Maximum-likelihood phylogenetic trees were constructed from alignments of these proteins (S1 Fig). In the resulting phylogeny, the Nif proteins of strain LKSZ1 clustered robustly within a monophyletic clade comprising Nif proteins derived from filamentous non-heterocystous cyanobacteria, including *Thermoleptolyngbya* sp. O-77 [16,17] and *Leptolyngbya valderiana* [18]. Support values for the relevant nodes exceeded, 90% for SH-like approximate likelihood ratio tests and 80% for ultrafast bootstrap across all analyzed proteins (S1 Fig), indicating a robust phylogenetic association. This phylogenetic placement was consistent among individual Nif proteins, demonstrating that the *nif* gene cluster of strain LKSZ1 is more closely related to those of filamentous cyanobacteria than to those of other unicellular diazotrophic cyanobacteria such as *Cyanothece* or *Crocosphaera* (Fig. 2 and S1 Fig).

To further examine the structural similarity of the *nif* gene cluster, the organization of the cluster in *Synechocystis* sp. LKSZ1 was compared with that of *Thermoleptolyngbya* sp. O-77, which showed the closest phylogenetic affinity in Nif protein analyses. Pairwise comparison using tBLASTx-based DiGAlign revealed extensive conservation of gene content and synteny between the two clusters (Fig. 6B). In both strains, the region spanning from *cnfR* to *mopII* exhibited a highly similar gene order and orientation, with strong amino acid-level similarity across corresponding genes. This conserved region included the core nitrogenase structural genes and genes involved in cofactor biosynthesis, indicating preservation of the overall functional architecture of the *nif* gene cluster. In contrast, structural differences were observed in the intervening region between genes encoding a *nifP* and NifX-associated nitrogen fixation protein, where an inversion was detected between the clusters of strain LKSZ1 and *Thermoleptolyngbya* sp. O-77 (Fig. 6B). Additionally, *modA*, *modBC,* and the gene located between them, as well as those positioned between *nifT* and *cnfR* in strain LKSZ1, were not shared with the corresponding genomic region in *Thermoleptolyngbya* sp. O-77. These results indicate that while the core *nif* architecture is conserved, local rearrangements and differences in flanking gene content exist.

To examine genomic features associated with the *nif* gene cluster in *Synechocystis* sp. LKSZ1, local nucleotide composition was analyzed across the chromosome (Fig. 5, Fig. 6B). Analysis of GC content revealed a pronounced decrease at both boundaries of the *nif* gene cluster. This reduction in GC content was restricted to the region encompassing the *nif* gene cluster and its immediate flanking sequences. In addition, analysis of GC skew showed localized deviations within the *nif* gene cluster, indicating a compositional divergence from surrounding chromosomal regions. Despite these compositional signatures, no insertion sequences or other identifiable mobile genetic elements were detected within or immediately adjacent to the *nif* gene cluster (Fig. 5).

### Genetic evidence for the retention of nitrogenase activity in *Synechocystis* sp. LKSZ1

Growth of *Synechocystis* sp. LKSZ1 under nitrogen-depleted conditions was examined on nitrogen-free BG11₀ solid medium under different light and oxygen regimes. As a non–nitrogen-fixing reference, *Synechocystis* sp. PCC 6803 was analyzed in parallel. Under anaerobic conditions, strain LKSZ1 exhibited clear growth on BG11₀ plates regardless of the light regime tested, including continuous illumination and a 12 h light/12 h dark cycle (Fig 7). In contrast, *Synechocystis* sp. PCC 6803 showed little to no growth under the same conditions. Strain LKSZ1 also displayed detectable growth under microoxic conditions combined with a light-dark cycle, whereas the growth of PCC 6803 remained severely impaired. Under fully aerobic conditions, growth of strain LKSZ1 on BG11₀ medium was limited, and no substantial differences were observed among the tested light regimes. These results demonstrate that *Synechocystis* sp. LKSZ1 is capable of sustained growth in the absence of combined nitrogen under low oxygen to anaerobic conditions, whereas the reference strain PCC 6803 is not.

**Figure 7.**
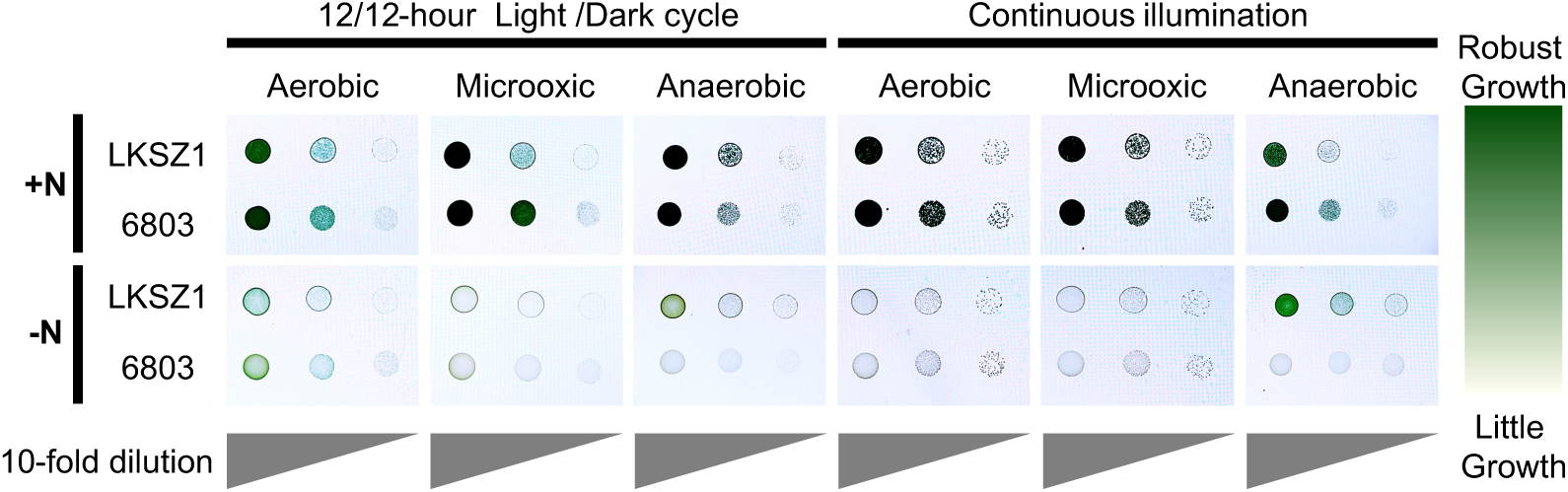
Growth of *Synechocystis* sp. LKSZ1 under nitrogen-depleted conditions. Spot growth assays of *Synechocystis* sp. LKSZ1 and *Synechocystis* sp. PCC 6803 on nitrogen-free BG11₀ agar under different oxygen and light conditions. Synchronized cells were washed with sterile water and adjusted to OD₇₅₀ = 0.2 prior to tenfold serial dilutions (10⁰, 10⁻¹, 10⁻²) and spotting (10 μL) onto plates. Plates were incubated under aerobic (ambient air), microoxic (6–12% O₂, 5–8% CO₂), or anaerobic (≤0.1% O₂, 10% CO₂) conditions. Light regimes consisted of either a 12-h light/12-h dark cycle (20 μmol photons m⁻² s⁻¹) or continuous illumination at 20 μmol photons m⁻² s⁻¹. Robust growth is indicated by dark green to green coloration resulting from dense cell accumulation, whereas growth inhibition is indicated by pale yellowish or nearly colorless spots with little or no visible biomass.

To determine whether the nitrogen-fixing phenotype observed in *Synechocystis* sp. LKSZ1 depends on the *nif* gene cluster, we attempted to isolate and characterize disruption mutants targeting *nifK*, which encodes the nitrogenase β-subunit. After introduction of the plasmid construct to disrupt *nifK*, two independent single cross-over recombinants (Δ*nifK*-1 and Δ*nifK*-2) could be isolated (Fig. 8). PCR analysis confirmed that *nifK* was disrupted in Δ*nifK*-1. Δ*nifK-2* still retains undisrupted *nifK*, although its native promoter was deleted. Under anaerobic conditions on nitrogen-free BG11₀ agar, the wild-type strain LKSZ1 exhibited robust growth, whereas both Δ*nifK*-1 and Δ*nifK*-2 showed little to no growth (Fig 9), indicating that *nifK* or its native promoter is required for photoautotrophic growth under nitrogen depleted conditions. As expected, *Synechocystis* sp. PCC 6803 did not grow under the same conditions. This growth defect in the mutants was consistently observed across replicate experiments.

**Figure 8.**
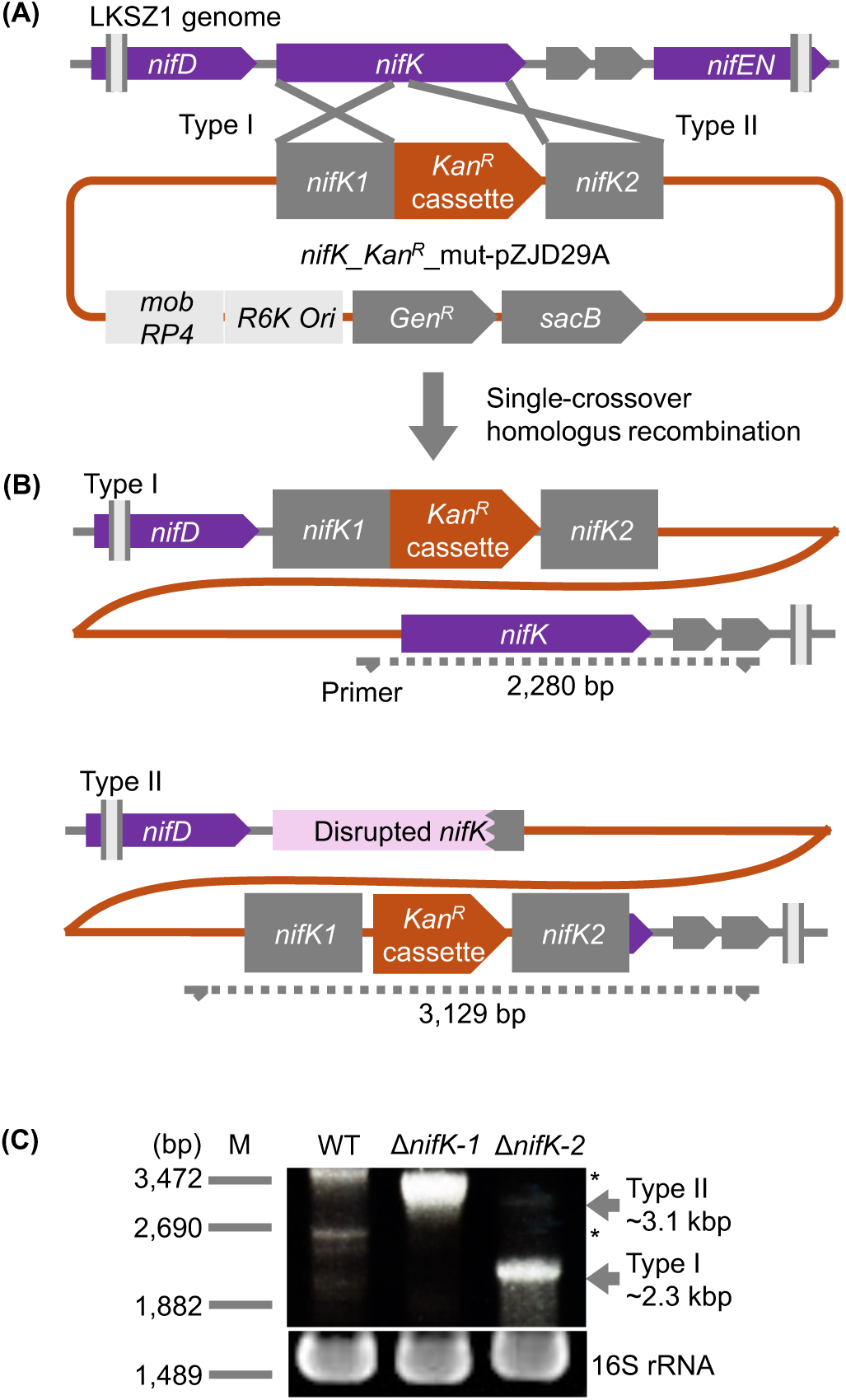
Construction and validation of *nifK* disruption mutants in *Synechocystis* sp. LKSZ1. (A) Schematic representation of the genomic *nifK* locus in strain LKSZ1 and the disruption construct used for homologous recombination. The suicide plasmid (pZJD29A derivative) contained upstream and downstream homologous regions flanking the *nifK* coding sequence and a kanamycin resistance cassette (*Kan^R^*) inserted between them. Homologous regions are indicated by gray boxes, and antibiotic resistance markers are shown in orange. (B) Proposed single-crossover recombination events leading to plasmid integration into the chromosomal *nifK* locus and the positions of diagnostic primers. Integration of the entire plasmid disrupts the *nifK* coding region. Two possible recombination configurations were observed: Type I (*nifK1* recombination) and Type II (*nifK2* recombination). Diagnostic PCR yields fragments of approximately 2.3 kbp for Type I and 3.1 kbp for Type II. (C) PCR verification of recombinant strains. Primer pairs spanning the plasmid backbone and chromosomal regions flanking the *nifK* locus were used to confirm insertion of the disruption construct. PCR products consistent with plasmid integration were detected in mutant strains but not in the wild-type control (asterisks indicate nonspecific bands). Δ*nifK*-1 corresponds to Type II recombination and Δ*nifK*-2 corresponds to Type I recombination. Amplification of the *16S rRNA* gene served as a positive control. These results demonstrate that the *nifK* disruption mutants were generated via single-crossover homologous recombination and confirm the genetic basis of the loss-of-function phenotype.

**Figure 9.**
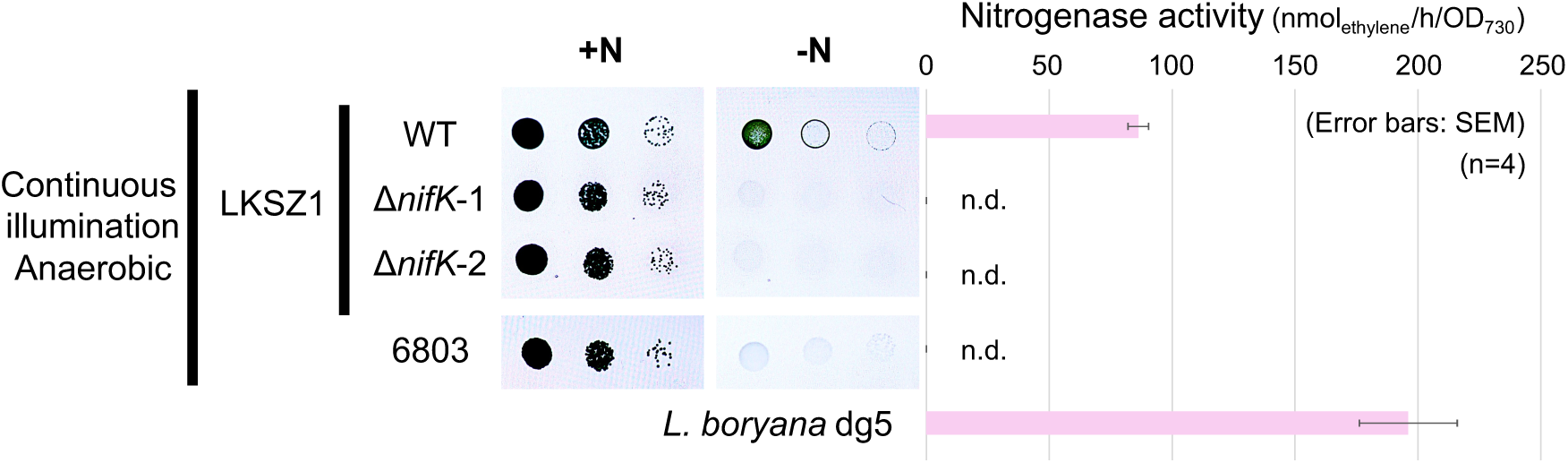
Genetic evidence for functional nitrogenase activity in *Synechocystis* sp. LKSZ1. Spot growth assays of wild-type (WT) strain LKSZ1, two independent *nifK* disruption mutants (Δ*nifK*-1 and Δ*nifK*-2), and *Synechocystis* sp. PCC 6803 on nitrogen-free BG11₀ agar under anaerobic conditions. Cells were adjusted to OD₇₅₀ = 0.2, serially diluted (10⁰, 10⁻¹, 10⁻²), and spotted onto plates. Under nitrogen-depleted anaerobic conditions, WT LKSZ1 exhibited robust growth, whereas both *nifK* disruption mutants failed to grow, comparable to PCC 6803. Acetylene reduction activity of *Synechocystis* sp. LKSZ1 (WT), two independent *nifK* disruption mutants (Δ*nifK*-1 and Δ*nifK*-2), *Synechocystis* sp. PCC 6803, and non-hererocystous nitrogen-fixing filamentous cyanobacterium *Leptolyngbya boryana* dg5 measured under anaerobic continuous illumination conditions. Cells were preconditioned under nitrogen-depleted conditions before the assay. Ethylene production was quantified by gas chromatography following incubation with acetylene and dithionite. Robust ethylene production was detected in WT LKSZ1 and *L. boryana* dg5, whereas no detectable activity was observed in either *nifK* disruption mutant or in PCC 6803. Data represent means ± SEM from independent biological replicates (n = 4). n.d., not detected.

Nitrogenase activity in *Synechocystis* sp. LKSZ1 was assessed by acetylene reduction assays under anaerobic conditions following growth in nitrogen-depleted medium. Robust ethylene production was detected in strain LKSZ1 (Fig 9), including in the absence of the external reductant dithionite (Fig. 10), indicating active nitrogenase function. In contrast, no ethylene production was observed in *Synechocystis* sp. PCC 6803 under the same conditions. Furthermore, both *nifK* disruption mutants failed to produce detectable ethylene under anaerobic conditions (Fig. 9). Collectively, these findings establish that nitrogenase activity and diazotrophic growth in strain LKSZ1 require an intact *nifK* gene and a functional *nif* gene cluster.

**Figure 10.**
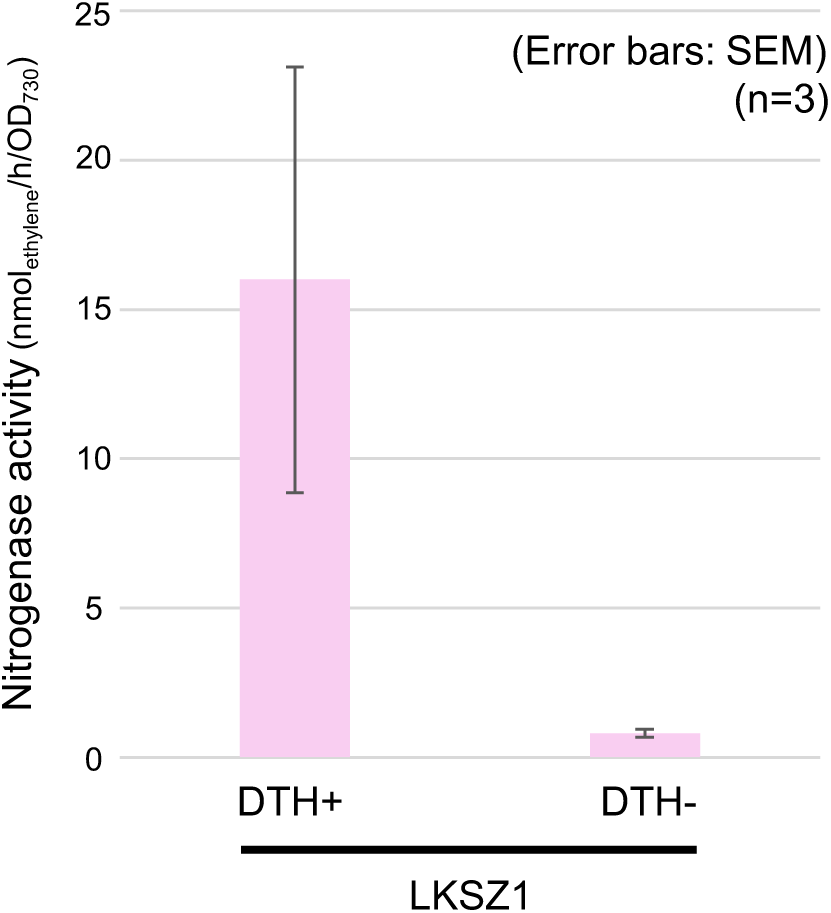
Influence of an exogenous reductant on nitrogenase activity in *Synechocystis* sp. LKSZ1. Acetylene reduction assay (ARA) of *Synechocystis* sp. LKSZ1 performed under anaerobic conditions in the presence or absence of the external reductant dithionite (DTH). Cells were preconditioned under nitrogen-depleted conditions before the assay. Ethylene production was quantified by gas chromatography following incubation with acetylene under strictly anaerobic conditions. Data represent means ±SEM from three independent biological replicates (n = 3). Ethylene production was detected in strain LKSZ1 even in the absence of exogenous DTH, indicating that the nitrogenase system is intrinsically functional and capable of utilizing endogenous reductants.

## Discussion

The *nif* gene cluster of strain LKSZ1 was most likely acquired via horizontal gene transfer, as supported by phylogenetic analyses (S1 Fig), *Synechocystis* pangenome comparisons (Fig. 4, Fig. 6), and sequence composition analyses (Fig. 5, Fig 6). Specifically, the *nif* gene cluster of strain LKSZ1 clusters with those of filamentous, non-heterocystous cyanobacteria of the *Thermoleptolyngbya/Leptolyngbya* lineage (S1 Fig), even though these taxa belong to different families (Fig. 2). The *nif* gene cluster of strain LKSZ1 is located within a region of genome plasticity enriched in shell and cloud genes and displays GC content and GC skew distinct from the surrounding chromosome (Fig. 6B). Such compositional signatures are characteristic of horizontally acquired DNA and indicate that the cluster was introduced as a discrete genomic unit [14,19], consistent with HGT. Although no mobile genetic elements were detected adjacent to the *nif* gene cluster (Fig. 5), this does not contradict the HGT hypothesis, as such elements may have been lost during subsequent adaptation of the cluster to the host genome.

Filamentous, non-heterocystous cyanobacteria such as *Thermoleptolyngbya* and *Leptolyngbya* are widespread in freshwater habitats and are often associated with benthic or biofilm lifestyles [20]. These ecological settings provide ample opportunities for physical proximity and genetic exchange among cyanobacteria [21]. Consistent with this view, multiple studies support horizontal transfer of *nif* gene clusters within the phylum *Cyanobacteria* [2,22]. For instance, transfer of nitrogenase lineages from *Synechococcales* to *Nostocales* has been reported [8], and the freshwater unicellular picocyanobacterium *Vulcanococcus limneticus* (formerly classified as *Synechococcus* sp. LL) acquired a *nif* gene cluster from distantly related cyanobacterial lineages through HGT [23], In addition, Deltaproteobacteria have been implicated as potential donors of *nif* gene clusters to cyanobacterial lineages [8,21].Thus, the observed transfer of a *nif* gene cluster from a filamentous donor to a unicellular *Synechocystis* lineage is consistent with known patterns of horizontal gene flow within cyanobacterial communities.

*Synechocystis* sp. LKSZ1 was isolated from a small freshwater pond where dense growth of aquatic vegetation is observed seasonally (Fig. 1A). Freshwater ecosystems frequently experience chronic or seasonal depletion of combined nitrogen. Specifically, small ponds, in particular, often become nitrogen limited during summer to autumn [24] or under low-rainfall conditions [25]. Intensive uptake and storage of combined nitrogen by aquatic plants and their associated microorganisms further exacerbate nitrogen depletion [26]. In addition, anoxia in bottom waters may promote nitrogen fixation in strain LKSZ1, which operates without specialized cellular differentiation such as heterocyst formation and instead maintains a unicellular, non-heterocystous mode of nitrogen fixation (Fig. 1B). In such environments, recurrent nitrogen limitation elevates access to atmospheric nitrogen as a key determinant of population persistence and competitiveness, and the capacity for anaerobic, photoautotrophic nitrogen fixation may therefore confer a decisive selective advantage. It should be noted that the microoxic and anaerobic conditions used in this study were generated using a gas-generating system that also elevates CO₂ concentrations. Therefore, the potential contribution of increased CO₂ levels to the observed growth phenotypes cannot be fully excluded.

However, acquisition of a *nif* gene cluster alone is not sufficient to establish nitrogen fixation. Indeed, although *V. limneticus* was suggested to have acquired a *nif* gene cluster through HGT, nitrogen fixation activity has not been experimentally confirmed in this organism [23]. More broadly, several cyanobacteria, including thermophilic *Synechococcus* [7,8,27], *Acaryochloris* [28], and *Cyanothece* [29] harbor *nif* gene clusters acquired through HGT. However, experimental evidence demonstrating sustained growth under nitrogen-depleted conditions remains lacking, to our knowledge.

Successful expression requires compatibility with the host’s metabolic, regulatory, and physiological framework [4–6,9,30,31]. The results of this study suggest that the genomic background of *Synechocystis* sp. LKSZ1 provided a permissive context that allowed functional integration of the horizontally acquired *nif* gene cluster. Indeed, extensive efforts to engineer nitrogen fixation into non-diazotrophic organisms, including *Synechocystis* sp. PCC 6803 and other heterologous hosts, have demonstrated that the simple introduction of *nif* genes is insufficient to confer activity. Reconstruction of nitrogenase function has required coordinated optimization of gene dosage, transcriptional regulation, oxygen protection mechanisms, and metal cofactor biosynthesis [32–36]. These studies highlight the substantial physiological barriers to functional *nif* expression. In contrast, strain LKSZ1 appears to have accommodated the *nif* gene cluster within a pre-existing metabolic and regulatory framework compatible with its energetic and redox constraints, underscoring the importance of host genomic permissiveness in the evolutionary stabilization of horizontally acquired metabolic traits.

Control of intracellular oxygen levels represents another critical constraint. Nitrogenase is highly oxygen sensitive [1–3,9], yet strain LKSZ1 exhibited nitrogenase activity under low-oxygen and anaerobic conditions, apparently without specialized nitrogen-fixing cells (Fig. 1B, Fig 7). Genes involved in low-oxygen tetrapyrrole biosynthesis and respiratory electron transport are conserved in the *Synechocystis* core genome (S2 File) and may contribute to maintaining intracellular redox conditions compatible with nitrogen fixation [8,30,37,38]. Such systems likely existed before *nif* acquisition and may have facilitated the functional deployment of the introduced cluster.

Anoxygenic photosynthesis components such as SQR appear to be important for nitrogen fixation [8]. Strain LKSZ1 retains genes associated with alternative, potentially anoxygenic electron transfer pathways, including *fccAB* (S2 File). In purple and green sulfur bacteria, FccAB functions as a flavocytochrome *c* sulfide dehydrogenase involved in anoxygenic photosynthesis [39]. Although the role of FccAB in *Synechocystis* remains unclear, its presence in strain LKSZ1 raises the possibility that alternative electron transfer routes may contribute to maintaining redox balance compatible with nitrogen fixation under low-oxygen conditions.

At the regulatory level, global nitrogen control systems, such as transcriptional factor NtcA [9,36,40] conserved in *Synechocystis,* coexist with cluster-encoded regulators such as CnfR [38,40]. This combination likely enables coordination between host nitrogen status and *nif* gene expression, allowing the horizontally acquired cluster to respond appropriately to environmental nitrogen availability. Such regulatory compatibility reduces the barrier for functional assimilation of foreign metabolic modules.

Results of this suggest that a combination of lateral gene donation, environmental nitrogen stress, and host metabolic flexibility drove the emergence of nitrogen fixation in a unicellular *Synechocystis* lineage via HGT. Testing this hypothesis will require comparative analyses of nitrogen-fixing cyanobacterial diversity in the Suzukake pond, quantitative monitoring of nitrogen dynamics, and genetic and physiological studies clarifying how strain LKSZ1 overcomes the oxygen paradox. These findings provide new insights into the gain and loss of nitrogen-fixing ability in cyanobacteria and may inform strategies aimed at conferring nitrogen fixation capacity to non-diazotrophic organisms, including crop plants.

## Materials and Methods

### Bacterial strains and culture conditions

*Synechocystis* sp. LKSZ1 was isolated from a freshwater pond located on the Suzukakedai campus of the Institute of Science Tokyo [15] and maintained as a clonal culture. *Synechocystis* sp. PCC 6803 was used as a non–nitrogen-fixing reference strain. *Leptolyngbya boryana* dg5 was used as a nitrogen-fixing reference strain. Unless otherwise stated, all strains were grown at 30-32 °C under continuous illumination (10-20 µmol photons m⁻² s⁻¹) in BG11 medium [41].

### Microscopy

Cell morphology of *Synechocystis* sp. LKSZ1 was examined using an all-in-one imaging system (APEXVIEW APX100, Evident/Olympus). Cells were collected from cultures grown under continuous light conditions, gently resuspended in fresh BG11 medium as needed, and mounted on glass slides for observation. Images were acquired using a 100× oil-immersion apochromatic objective lens (numerical aperture 1.45). Representative micrographs were processed using the manufacturer’s software without altering relative contrast or brightness.

### ANI calculation, Genome annotation, and Pangenome analysis

To characterize genomic diversity within the genus *Synechocystis* and to determine the genomic divergence of strain LKSZ1, comparative genomic analyses were performed using publicly available genome sequences. A total of 32 *Synechocystis* genomes deposited in the RefSeq database of the National Center for Biotechnology Information [42] were analyzed, including complete genomes, chromosomes, scaffolds, and contig-level assemblies. The chromosome sequence of *Synechocystis* sp. LKSZ1 (RefSeq accession: NZ_AP031572) was included in this dataset.

Average nucleotide identity (ANI) values were calculated using the ANIb algorithm implemented in pyani-plus (v1.0.0). Genomes were fragmented into 1,020-bp segments prior to pairwise BLASTn comparisons following default parameters. The resulting similarity matrix was subjected to hierarchical clustering to assess genome-wide relatedness.

To ensure consistency across genomes, all sequences were uniformly reannotated using Bakta (v1.11.3) with the Bakta database v6.0 (DOI: 10.5281/zenodo.14916843) [43]. Bakta annotations were used as input for downstream analyses. Pangenome reconstruction and analysis were performed using PPanGGOLiN (v2.2.4) [13] with the “all” workflow under identity threshold 0.5 and coverage threshold 0.5 parameters. Gene families were classified into persistent, shell, and cloud categories based on their frequency across genomes based on the statistical partitioning model implemented in PPanGGOLiN, and the relative proportions of each category were calculated for each genome. Based on this classification, PPanGGOLiN further identified regions of genome plasticity (RGPs) as genomic segments statistically enriched in shell and cloud gene families that interrupt stretches dominated by persistent genes [14]. These regions were detected automatically by PPanGGOLiN according to its internal criteria and were interpreted as genomic regions with an increased likelihood of gene gain, loss, or rearrangement, frequently associated with horizontal gene transfer.

### Identification of the *nif* gene cluster

Identification of the nitrogen fixation gene cluster in *Synechocystis* sp. LKSZ1 was based on the results of the pangenome analysis and prior definitions of cyanobacterial *nif* gene organization. In the chromosome sequence of strain LKSZ1 (RefSeq accession: NZ_AP031572), a contiguous genomic region spanning locus tags ABXS88_RS01395 to ABXS88_RS01550 was infered as complete *nif* and *nif* related gene encoded region. Following established criteria used in previous studies [33–36,40], the boundaries of the *nif* gene cluster were defined as extending from the molybdate transporter gene fusion *modBC* to a gene annotated as a putative transcriptional regulator (*cnfR*) associated with *nif* gene expression. This region encompassed core nitrogenase structural genes (*nifHDK*), genes required for cofactor biosynthesis (*nifENB*), and additional accessory genes typically associated with functional nitrogen fixation systems.

### Phylogenetic analyses

Phylogenetic analyses were conducted to determine the evolutionary position of *Synechocystis* sp. LKSZ1 within Cyanobacteria and to infer the evolutionary origin of its *nif* gene cluster. For genome-wide phylogenetic placement, a phylogeny of species representative based on 120 conserved bacterial marker genes was obtained from the Genome Taxonomy Database (GTDB) [44].Based on the latest R226 dataset, cyanobacterial lineages were extracted and visualized on iTOL v7 web server [45].

To compare the species phylogeny of cyanobacteria with the the phylogeny of their *nif* gene cluster, amino acid sequences of key nitrogenase components and cofactor biosynthesis proteins (NifH, NifD, NifK, and NifEN) were selected. A subset of Chen’s dataset (comprising 650 cyanobacterial lineages) was used as a reference [8], and the corresponding sequences from strain LKSZ1 were added. Only taxa encoding full set of target proteins were retained for analysis. Outgroup genes were chosen according to previous studies, using BchL/ChlL as the outgroup for NifH, BchB/ChlB for NifD, BchN/ChlN for NifK, and BchNB/ChlNB for NifEN [46].

Each sequence group was multiple aligned using MAFFT (v7.526) with the L-INS-i algorithm [47,48]. Poorly aligned regions were removed using trimAl (v1.4) with the gappyout option [49]. For taxa in which NifE and NifN were encoded as separate proteins, the two sequences were concatenated to match the fused NifEN protein encoded in strain LKSZ1. The trimmed alignments were used to infer maximum-likelihood phylogenetic trees with IQ-TREE 2 (v2.3.6) [50]. The best-fitting amino acid substitution model was selected automatically using ModelFinder based on the Bayesian information criterion. Branch support was assessed using 1,000 ultrafast bootstrap replicates and SH-like approximate likelihood ratio tests.

### Comparative analysis of the *nif* gene cluster and genomic signatures of HGT

To examine the evolutionary origin of the *nif* gene cluster in *Synechocystis* sp. LKSZ1, comparative analyses of gene cluster organization and genomic compositional features were performed. For structural comparison, the *nif* gene cluster of *Thermoleptolyngbya* sp. O-77, which showed closely related to the NifDKEN proteins of strain LKSZ1, was selected.

60 kbp genomic regions encompassing the *nif* gene clusters were extracted from the chromosome sequences of strain LKSZ1 and *Thermoleptolyngbya* sp. O-77 (RefSeq accession: NZ_AP017367.1). Structural similarity and synteny between the two regions were assessed using DiGAlign (v2.0) with tBLASTx as the homology search algorithm [51]. Nucleotide sequences were translated in six frames, and protein-level synteny was evaluated to visualize conserved gene order, orientation, and rearrangements.

To evaluate genomic signatures associated with a presumed HGT event, GC content and GC skew were calculated for the *nif* gene cluster and its flanking regions using Proksee web server [52]. A sliding window size of 10,000 bp with a step size of 100 bp was applied to detect local deviations from genome-wide compositional patterns. These metrics were visualized on circular genome maps to assess compositional discontinuities associated with the *nif* gene cluster.

In addition, the presence of mobile genetic elements in the *nif* gene cluster and its surrounding region was examined using the mobileOG-db integrated in Proksee web server [53]. These analyses were used to identify genomic features commonly associated with horizontally acquired gene clusters.

### Construction of *nifK* disruption mutants

To determine whether the nitrogen-fixing phenotype of *Synechocystis* sp. LKSZ1 depends on the *nif* gene cluster; a disruption mutant targeting the nitrogenase β-subunit gene *nifK* was constructed by homologous recombination using a conjugation-based DNA transfer system. Primers that were used are shown in S1 Table.

For the construction of the disruption plasmid, approximately 700-bp upstream (698 bp, Primers 004 and 012) and downstream (697 bp, Primers 005 and 015) homologous regions, including the *nifK,* were amplified by PCR using KOD One DNA Polymerase (TOYOBO, Japan). A kanamycin resistance cassette driven by the *pc* promoter was amplified from Ppc_Kan-pUC18 (Primers 013 and 014), which was derived from kanamycin resistance gene in S6803-alphaY252C [54] and pc promoter in Tp/pUC18. Tp/pUC18 is a derivative of pUC18 [55] in which the *pc* promoter and a trimethoprim resistance gene were inserted into the multiple cloning site; the resistance cassette can be readily exchanged via digestion with PshAI and XhoI, allowing flexible replacement of selectable markers. The homologous arms and the kanamycin resistance cassette were assembled into pUC19 using the In-Fusion HD Cloning Kit (TaKaRa Bio, Japan) and propagated in *Escherichia coli* DH5α. The resulting recombinant fragment was re-amplified by PCR (Primers 022 and 023) and inserted into the suicide vector pZJD29A [56] by In-Fusion cloning.

The final construct was propagated in *E. coli* JM109 λ/pir and acquired a plasmid *nifK*_*Kan^R^*_mut-pZJD29A. The plasmid introduced into *E. coli* S17-1 λ/pir by electroporation for use as the donor strain. Biparental conjugation was performed by mixing donor cells with exponentially growing LKSZ1 cells (OD_750_ ≈ 0.64), spotting the mixture onto BG11 agar plates, and incubating for 2 days at 32 °C under continuous light.

Following conjugation, cells were recovered and plated onto BG11 agar containing 5 µg/mL kanamycin, 20 µg/mL tetracycline, and 1% (w/v) sucrose. Cells were spread onto 0.45-µm PVDF membrane filters placed on selective plates and incubated at 32 °C under continuous illumination for approximately 3 weeks until resistant colonies appeared. To promote complete segregation of mutant alleles, colonies were repeatedly subcultured on BG11 plates containing 20 µg/mL kanamycin until stable transformants were obtained.

Successful recombination at the *nifK* locus was verified by diagnostic PCR. Genomic DNA was extracted from wild-type and putative mutant strains using phenol–chloroform extraction [15]. PCR reactions were performed using LA Taq DNA Polymerase (TaKaRa Bio, Japan) according to the manufacturer’s instructions. To confirm plasmid integration via single-crossover homologous recombination, primer 011 (annealing within the pZJD29A plasmid backbone) was paired with primer 048 (annealing within the chromosomal region encoding the NifX-associated nitrogen fixation protein adjacent to the *nifK* locus). Amplification of the expected fragment was detected in mutant strains but not in the wild-type control, indicating integration of the suicide plasmid into the chromosomal *nifK* locus. As an internal control for DNA quality, the *16S rRNA* gene was amplified using universal primers 27f and 1525r. PCR products were analyzed by agarose gel electrophoresis and visualized under UV illumination.

### Growth assays under nitrogen-depleted conditions

Growth of *Synechocystis* sp. LKSZ1 wildtype (WT) and *nifK* mutant under nitrogen-depleted conditions were assessed using solid media assays. As a non–nitrogen-fixing control, *Synechocystis* sp. PCC 6803 was analyzed in parallel. Cultures were pre-grown under continuous light conditions and subsequently synchronized by dark incubation for 3 days, followed by a 12 h light/12 h dark cycle for an additional 3 days.

Cells were harvested by centrifugation, washed three times with sterile water to remove residual combined nitrogen, and resuspended to an optical density at 750 nm (OD₇₅₀) of 0.2. Tenfold serial dilutions (10⁰, 10⁻¹, and 10⁻²) were prepared, and 10 µL aliquots of each dilution were spotted onto BG11₀ agar plates lacking combined nitrogen [57]. Control plates added KNO_3_ (final concentration 15 mM) to BG11_0_ agar plates were prepared in parallel [33].

Spotted plates were incubated under different combinations of light and oxygen conditions. Light regimes included continuous illumination (20 µmol photons m⁻² s⁻¹) or a 12 h light/12 h dark cycle. Oxygen conditions were set as aerobic (ambient air), microoxic (6–12% O₂, 5–8% CO₂), or anaerobic (≤0.1% O₂, 10% CO₂). Microoxic and anaerobic environments were established using sealed incubation jars (Sugiyamagen, Japan) and commercially available gas-generating systems (Mitsubishi Gas Chemical).

Anaerobic conditions were verified using an oxygen indicator (Sugiyamagen, Japan), confirming that O₂ levels did not exceed 0.1% throughout incubation. In this system, residual and photosynthetically produced oxygen is continuously scavenged by the AnaeroPack reagents, although precise control or measurement of oxygen dynamics during illumination is not possible. The elevated CO₂ levels under microaerobic and anaerobic conditions are an inherent feature of the gas-generating system.

The *nifK* mutants were experimented with only anaerobic and continuous light conditions. All growth assays were conducted at 32 °C, and colony formation was monitored over time.

### Assay of nitrogenase activity

The nitrogenase activity of LKSZ1 was measured as acetylene reduction activity. For the induction of nitrogenase, the cells grown on BG-11 agar plate under aerobic conditions for 10-14 d were suspended in water (500 μL of OD_730_ = 10.0). The cell suspension was spread on nitrogen-free BG-11_0_ agar plate, and the plate was incubated in an anaerobic jar under continuous light at 30°C for 20 h. The cells on the plate were suspended in 1.0 mL of BG-11_0_ liquid medium in the anaerobic chamber. Aliquots (1 mL) of the cell suspensions were transferred into 8-mL glass vials (Cat No. V-5A, NICHIDEN RIKA GLASS Co., Ltd, Kobe). To detect stable nitrogenase activity, 0.5 M dithionite dissolved in 0.2 M NaHCO₃ solution was added to the suspension as needed (final concentration: 5 mM). Glass vials were enclosed with an air-tight septum and the gas phase of the vial was replaced by sparging with acetylene gas mixture (10% acetylene-90% in argon) for 10 sec. The vial was incubated under continuous light at 30°C for 30 min (Fig. 9) or 1 h (Fig. 10). A portion of the gas phase (500 µl) was analyzed by gas chromatography as described previously [38].

## Acknowledgments

We thank Akito Machida (Institute of Science Tokyo) for providing the photograph of Suzukake Pond. We are also grateful to the Integrative Bioscience Facility at the Institute of Science Tokyo, for technical assistance with microscopy analysis. This work is supported in part by MEXT/JSPS KAKENHI (22K06276 to SM) and COI-NEXT (JPMJPF2102 to YF) of the Japan Science and Technology Agency.

## Notes

### Competing Interest Statement

The authors have declared no competing interest.

