## Supplemental Figure and Table for "Horizontal gene transfer drives the emergence of nitrogen fixation in a unicellular *Synechocystis* lineage"

### Supporting information

(A)

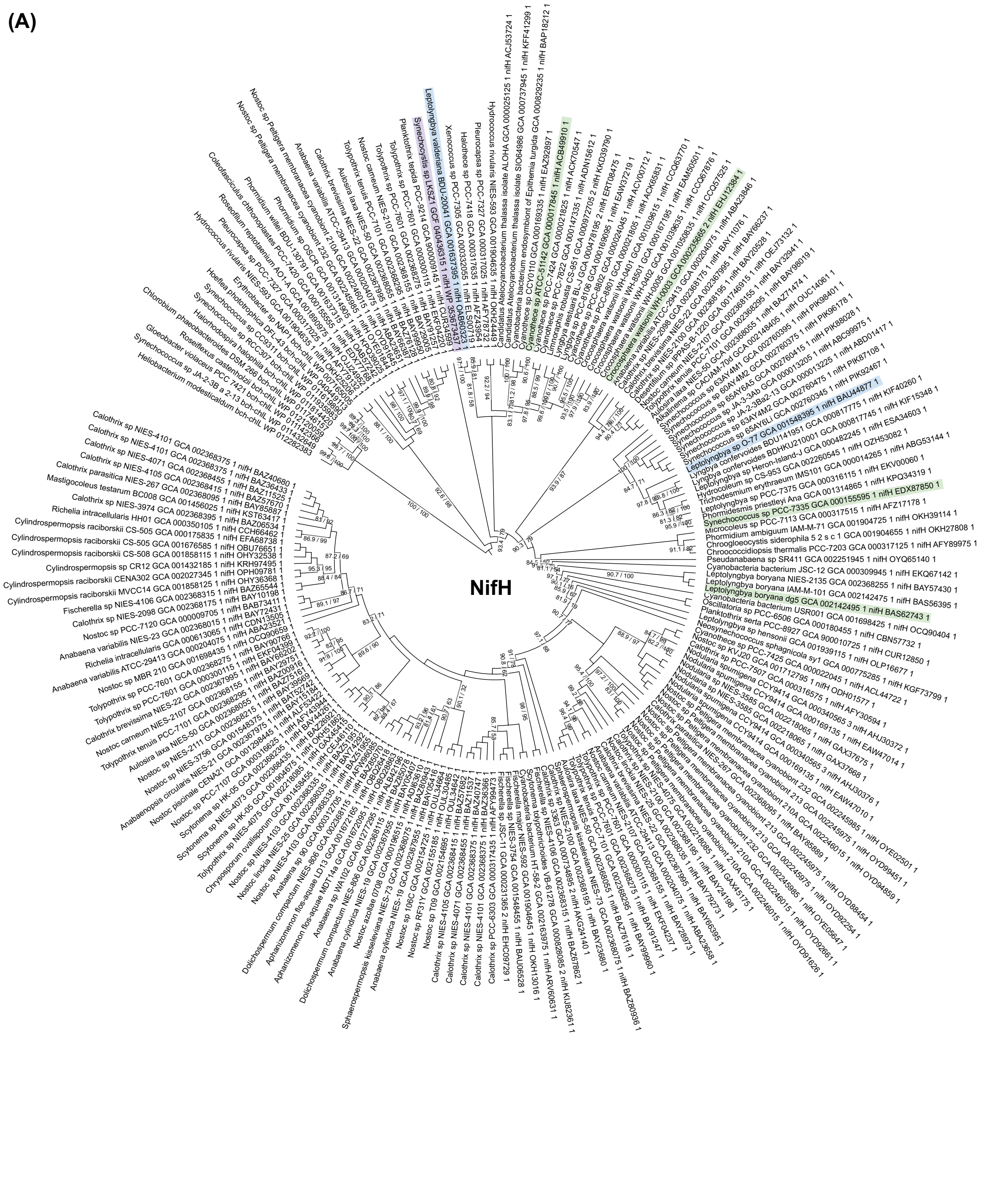

(B)

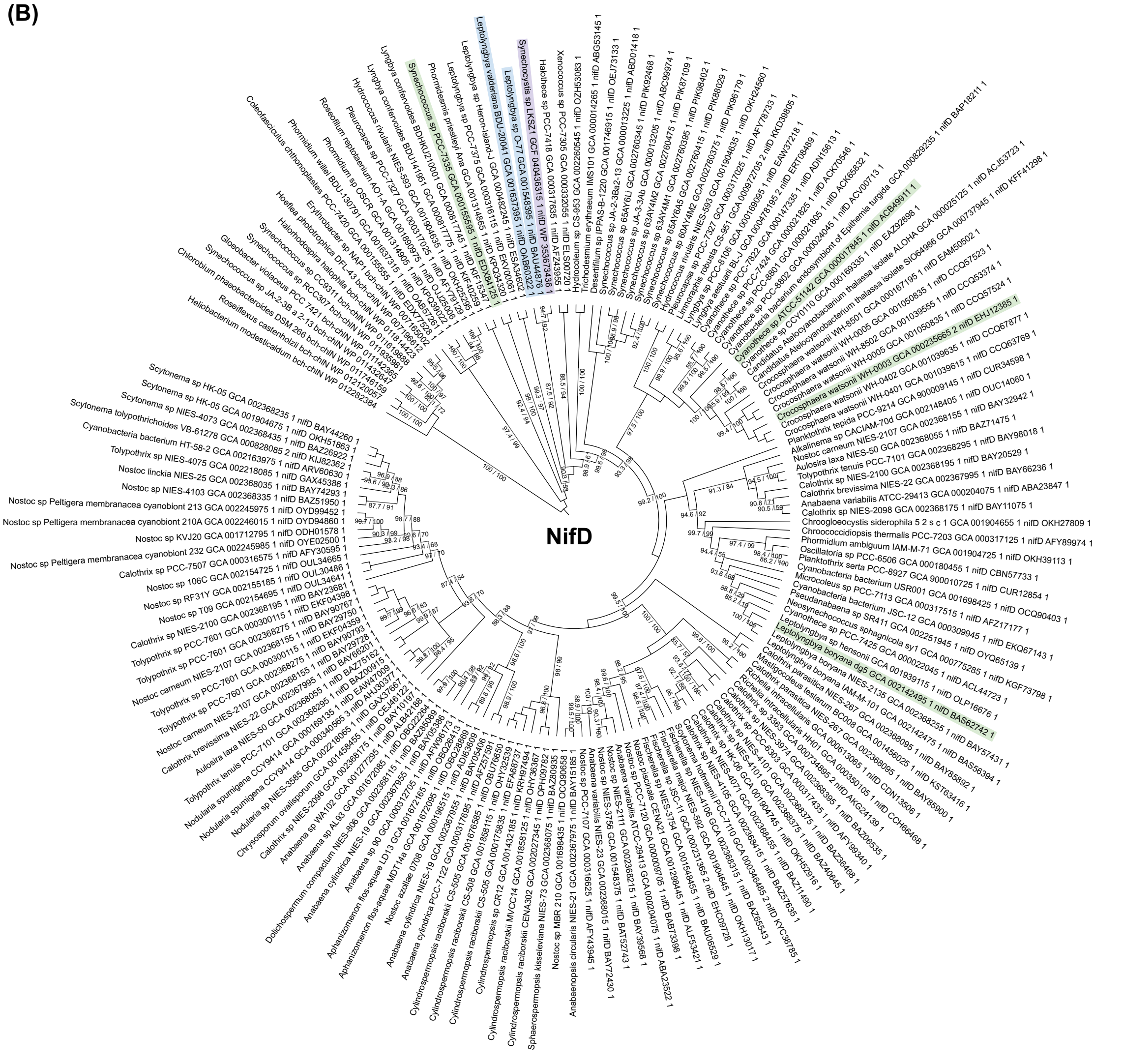


3

**(C)**

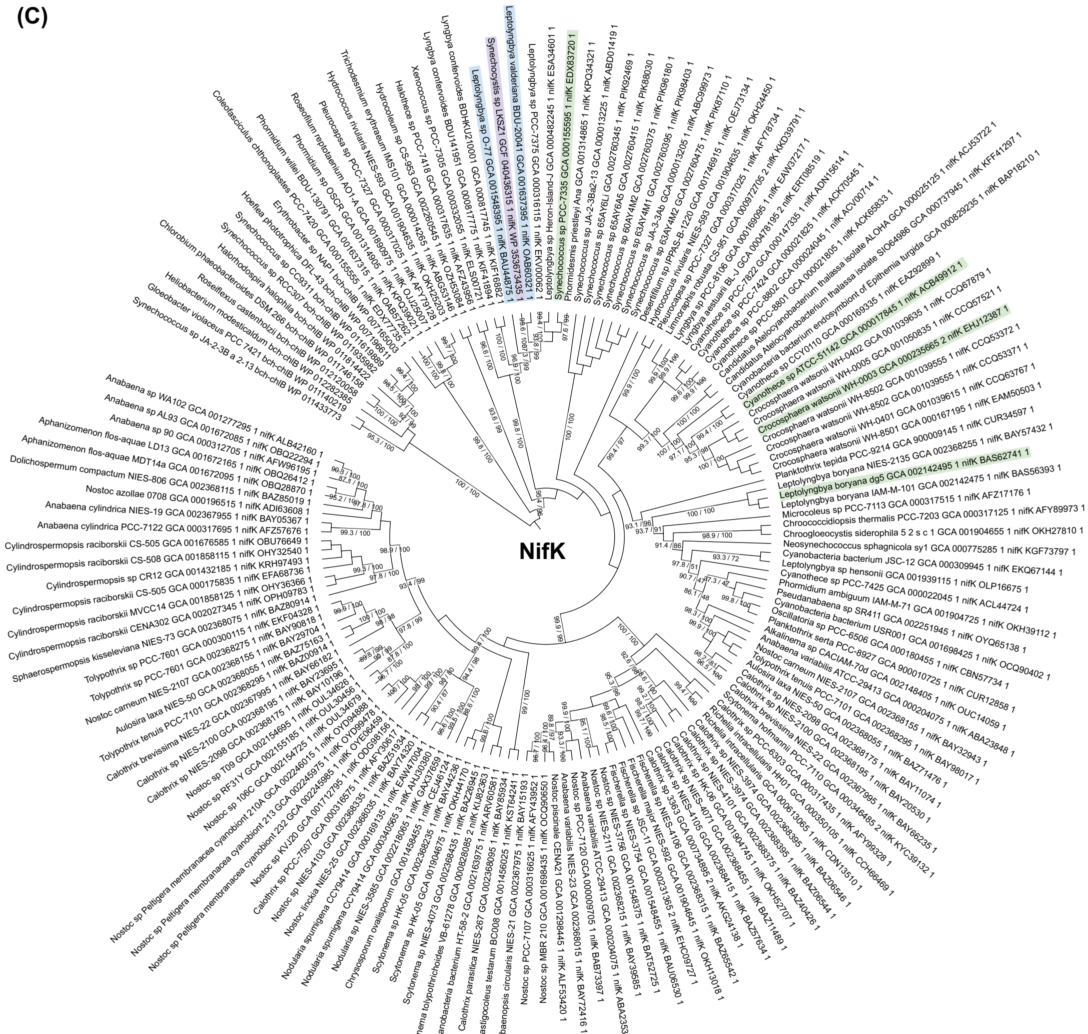


4

(D)

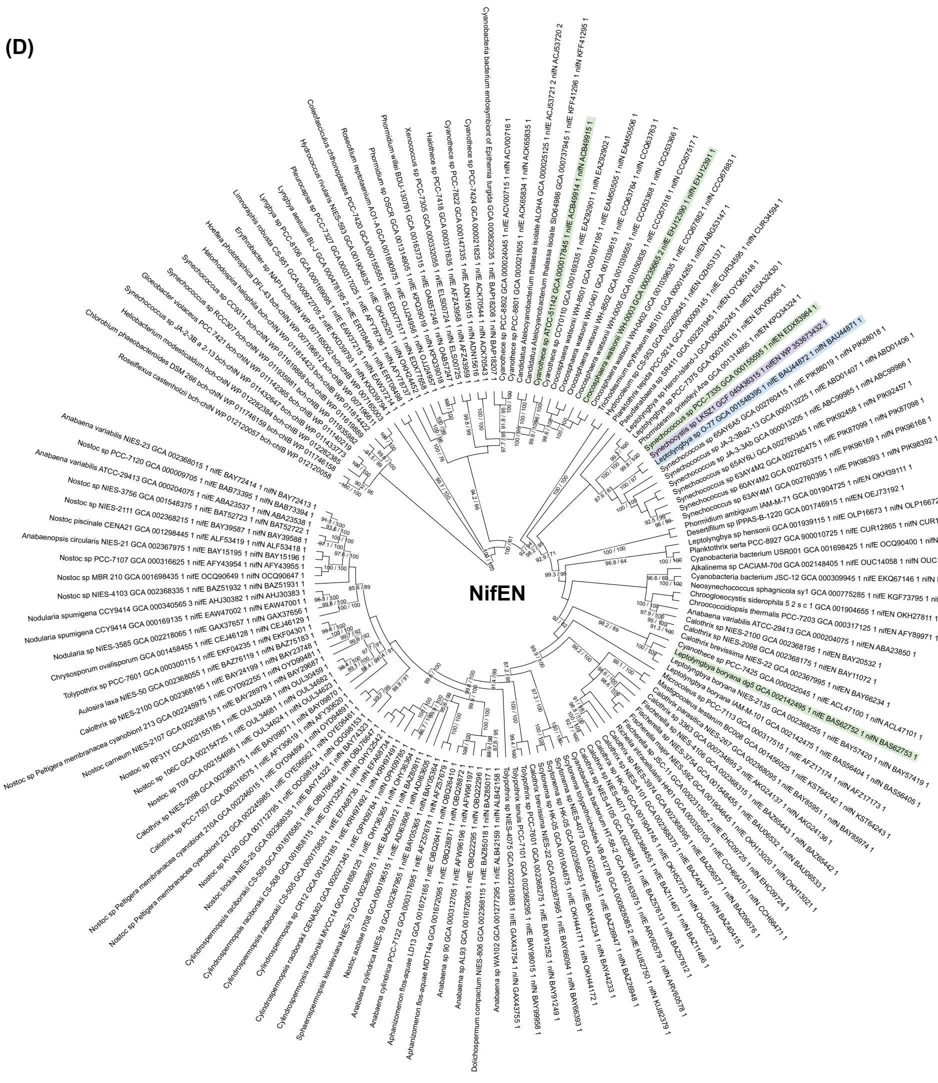


**S1 Fig. Maximum-likelihood phylogenetic trees of core nitrogenase proteins.** The amino acid sequences of (A) NifH, (B) NifD, (C) NifK, and (D) concatenated NifEN from representative cyanobacteria and related taxa was used. Sequences from *Synechocystis* sp. LKSZ1 were integrated into a curated dataset derived from previous reports (Chen et al., 2022; Pi et al., 2022). In panel D, for taxa in which NifE and NifN occur as separate proteins, sequences were concatenated in the order NifE–NifN to match the fused NifEN protein encoded in strain LKSZ1. Homologous bacteriochlorophyll biosynthesis proteins (Bch/ChlL, Bch/ChlN, Bch/ChlB, and Bch/ChlNB, respectively) were used as outgroups. Node support values represent SH-like approximate likelihood ratio tests and ultrafast bootstrap (1,000 replicates) (SH-aLRT/UFBoot).

Nif proteins from strain LKSZ1 (highlighted in purple) consistently cluster with those from filamentous non-heterocystous cyanobacteria (blue), including *Thermoleptolyngbya* sp. O-77 (formerly *Leptolyngbya* sp. O-77) and *Leptolyngbya valderiana*, rather than with unicellular diazotrophic genera (green) such as *Cyanothece* sp. ATCC 51142 (synonymous with *Crocosphaera subtropica* ATCC 51142) or *Crocosphaera watsonii*. *Synechococcus* sp. PCC 7335 and *Leptolyngbya boryana* dg5 are also highlighted in green. This phylogenetic placement strongly suggests a horizontally acquisition of the *nif* gene cluster in strain LKSZ1 from a phylogenetically distinct cyanobacterial clade.

**S1 Table. Primer list**

| | |
|---|---|
| 004 | CGACTCTAGAGGATCCTTCCCTGGAACCGATCAT |
| 005 | CGGTACCCGGGGATCGGCGATCAAAAATGGGATAA |
| 011 | GGCTCGTATGTTGTGTG |
| 012 | GGTCGTGCGAACTGCTTTCAGCCTTTTTGCC |
| 013 | TGAAAGCAGTTCGCACGACCCAGTTGAC |
| 014 | GTTCCCGAATCAGCTCATTTCAGAATATTTGCTCG |
| 015 | AAATGAGCTGATTCGGGAACTGAAGCGAATTTTAG |
| 022 | GTTTTCCCAGTCACGACGTTGTAAAAC |
| 023 | CAGGAAACAGCTATGACCATGATTAC |
| 048 | GTTCAAAGGGGCAACAGCAG |
| 27f | AGAGTTTGATCCTGGCTCAG |
| 1525r | AAAGGAGGTGATCCAGCC |

**S1 File. Genome-level statistics for *Synechocystis* strains included in the PPanGGOLiN pangenome analysis.** Rows correspond to individual genomes identified by strain name and NCBI RefSeq assembly accession number. Columns report assembly metrics (number of contigs, predicted genes, fragmented genes), gene family statistics (total families, families with fragments, multicopy families, soft-core and exact-core families and genes), and partitioned persistent, shell, and cloud components (gene counts, family counts, and relative proportions), including fragmented and multicopy families. Quality metrics (completeness, contamination, fragmentation) and counts of regions of genome plasticity (RGPs), spots, and modules as defined by PPanGGOLiN are also provided (Gautreau et al., 2020; Bazin et al., 2020).

**S2 File. Gene family presence–absence matrix for the *Synechocystis* pangenome.** This matrix was generated using the PPanGGOLiN framework (Gautreau et al., 2020). Rows correspond to individual gene families and include comprehensive functional annotations and clustering statistics, such as gene identifiers, non-unique gene names, functional descriptions, number of isolates possessing the family, total number of sequences, average sequences per isolate, fragment classification, genomic order information, quality control metrics, and nucleotide group size statistics. Columns represent individual *Synechocystis* genomes (identified by strain name and NCBI RefSeq assembly accession number), with matrix entries indicating the presence or absence of each gene family across the dataset.
